# Quiescent cells maintain active degradation-mediated protein quality control requiring proteasome, autophagy and nucleus-vacuole junctions

**DOI:** 10.1101/2024.06.29.601343

**Authors:** Dina Franić, Mihaela Pravica, Klara Zubčić, Shawna Miles, Antonio Bedalov, Mirta Boban

## Abstract

Many cells spend a major part of their life in quiescence, a reversible state characterized by a distinct cellular organization and metabolism. In glucose-depleted quiescent yeast cells, there is a metabolic shift from glycolysis to mitochondrial respiration, and a large fraction of proteasomes are reorganized into cytoplasmic granules containing disassembled particles. Given these changes, the operation of protein quality control (PQC) in quiescent cells, in particular the reliance on degradation-mediated PQC and the specific pathways involved, remains unclear. By examining model misfolded proteins expressed in glucose-depleted quiescent yeast cells, we found that misfolded proteins are targeted for selective degradation requiring functional 26S proteasomes. This indicates that a significant pool of proteasomes remains active in degrading quality control substrates. Misfolded proteins were degraded in a manner dependent on the E3 ubiquitin ligases Ubr1 and San1, with Ubr1 playing a dominant role. In contrast to exponentially growing cells, the efficient clearance of certain misfolded proteins additionally required intact nucleus-vacuole junctions (NVJ) and Cue5-independent selective autophagy. Our findings suggest that proteasome activity, autophagy, and NVJ-dependent degradation operate in parallel. Together the data demonstrate that quiescent cells maintain active PQC that relies primarily on selective protein degradation. The necessity of multiple degradation pathways for the removal of misfolded proteins during quiescence underscores the importance of misfolded protein clearance in this cellular state.

## Introduction

Misfolded proteins can interfere with essential cellular processes and give rise to disease, such as Alzheimer’s, Parkinson’s and other (1–5). The protein quality control (PQC) system, a complex network of evolutionarily conserved pathways, prevents the accumulation of misfolded proteins by mediating protein refolding, selective degradation, and spatial sequestration to inclusions (1, 4, 6–8).

Terminally misfolded proteins are posttranslationally modified by polyubiquitin and subsequently targeted for degradation to the proteasomes. Protein ubiquitination is catalyzed by a series of enzymes, including ubiquitin-conjugating enzymes (E2) and ubiquitin protein ligases (E3) (9). The primary determinants of ubiquitination specificity are the E3 ubiquitin ligases, in many cases in cooperation with molecular chaperones (10). In yeast, the principal E3 ubiquitin ligases that target cytoplasmic misfolded proteins for degradation are Ubr1, a protein found in the cytoplasm and nucleus, a nuclear E3 ligase San1 and Doa10, the integral membrane E3 ligase of the endoplasmic reticulum and the inner nuclear membrane (11–18). The selectivity of proteasome degradation is achieved by the ubiquitin-binding receptors present in the 19S regulatory particle, while the substrates are cleaved by the proteolytic enzymes present within the 20S core particle (19). The 19S particles also contain ATPases that facilitate substrate unfolding and entry into the 20S core particle. In addition, 19S particles contain deubiquitinating enzymes that remove ubiquitin from the substrate prior to its entry into the 20S core particle, thereby facilitating substrate degradation (20).

Under the conditions of misfolded protein overload, such as in the case of proteasome inhibition or non-functional protein ubiquitination, cells can sequester misfolded proteins into subcellular deposition sites called INQ (intranuclear quality control compartment), JUNQ (juxtanuclear quality control compartment) and CytoQ (cytoplasmic quality control compartment) (21–26). These inclusions are dynamic (21, 22), and from there, misfolded proteins can be disaggregated and directed to refolding or degradation (27). In contrast, cytoplasmic insoluble protein deposits (IPOD) are cytoplasmic inclusions that terminally sequester insoluble misfolded proteins with the amyloid-like properties (8, 21).

Due to the structural restraints of the translocation channel, only individual proteins that have been unfolded by the ATPases in the 19S regulatory particle can be degraded by the proteasome (20), while larger assemblies or protein aggregates can be degraded by autophagy (28). In the process of autophagy, cargo material is enclosed by autophagosomes, double-membrane vesicles decorated with the ubiquitin-like protein Atg8, and delivered to the lysosomal compartment, or vacuole in yeast (29). The formation of autophagosomes is mediated by a set of conserved proteins, collectively termed the core machinery. In non-selective or bulk autophagy, autophagosome biogenesis is triggered by nutrient starvation, including glucose depletion (29). In selective autophagy, autophagosome biogenesis can be triggered even in the absence of starvation, by cargo receptors or autophagy receptors, which simultaneously bind cargo marked by a specific modification, such as ubiquitin, and Atg8 present on the autophagosomal membrane (30).

In nature many cells spend a considerable amount of their lifetime in a reversible non-dividing state known as quiescence, which is characterized by a distinct cellular organization and metabolism (31–34) When incubated in a rich glucose-based medium, yeast *Saccharomyces cerevisiae* grows exponentially until glucose is exhausted. At that point, the culture undergoes a diauxic shift, cells switch to utilizing non-fermentable carbon source, such as ethanol, and enter into quiescence (31, 35). Finally, upon exhaustion of the carbon source, cells cease dividing entirely, and the culture enters the stationary phase.

In contrast to dividing yeast cells, in which the majority of the proteasomes accumulate in the nucleus (36), in quiescent cells, proteasomes relocalize to the nuclear periphery and to cytoplasmic condensates called proteasome storage granules (37), which are thought to contain inactive proteasomes disassembled into core and regulatory particles (38, 39). How quiescent cells manage protein quality control substrates has remained unclear. Previous reports using stationary phase cultures have indicated the accumulation of misfolded proteins within inclusions (40) and a decline in the degradation of proteasomal substrates (39, 41). However, due to the barely detectable protein expression levels observed (39), the interpretation of the results was challenging.

In this study we investigated whether quiescent cells retain degradation-mediated protein quality control, by expressing model misfolded proteins in quiescent cells of yeast *Saccharomyces cerevisiae*. We report that quiescent cells target misfolded proteins for selective degradation by the proteasome, suggesting that a significant pool of the 26S proteasomes remain actively engaged in the degradation of quality control substrates during cell quiescence. Moreover, the efficient elimination of certain misfolded proteins was additionally dependent on the intact nucleus-vacuole junctions (NVJ) and core autophagy machinery. The requirement for the functional autophagy was substrate-specific, indicating selectivity, and was independent of the known yeast ubiquitin-binding cargo receptor Cue5. Together our results indicate that degradation-mediated PQC is sustained during cell quiescence and chronological aging.

## Results

### Glucose-depleted quiescent yeast cells retain degradation-mediated protein quality control

To investigate whether quiescent cells retain degradation-mediated protein quality control, we analyzed cytoplasmic misfolded proteins tGnd1 (truncated Gnd1) and stGnd1 (small truncated Gnd1), C-terminally truncated versions of yeast 6-phosphogluconate dehydrogenase enzyme Gnd1, that arise due to a premature stop codon in *GND1* gene (11). We expressed HA-epitope tagged tGnd1 and stGnd1 in quiescent yeast cells and examined protein stability by cycloheximide chase experiment (Fig. 1). Genes encoding s/tGnd1-HA were placed under the control of the constitutive *PIR3* promoter, which becomes active upon cell entry into quiescence (Fig. S1). Additionally, we analyzed exponentially growing cells that expressed s/tGnd1-HA from the constitutive *TEF1*-gene promoter. Cell cultures grown under identical conditions were tested for cell density, glucose and ethanol concentration at different time points, showing that by the time point of 24 hours after culture inoculation, cells have consumed all glucose (Fig. 1A). Ethanol was consumed by day five, which coincided with the complete cessation of cell growth, indicating cell culture entry into the stationary phase. In contrast to the wild type protein Gnd1, which was stable, model misfolded proteins tGnd1-HA and stGnd1-HA expressed in cells from two-day-old culture were unstable, indicating selective degradation of misfolded proteins, similarly as in exponentially growing cells (Fig. 1B). Misfolded proteins s/tGnd1 were also selectively degraded in a yeast strain of a different genetic background, W303 (Fig. 1C). Together the data indicate that glucose depleted quiescent yeast cells maintain degradation-mediated protein quality control.

**Figure 1.**
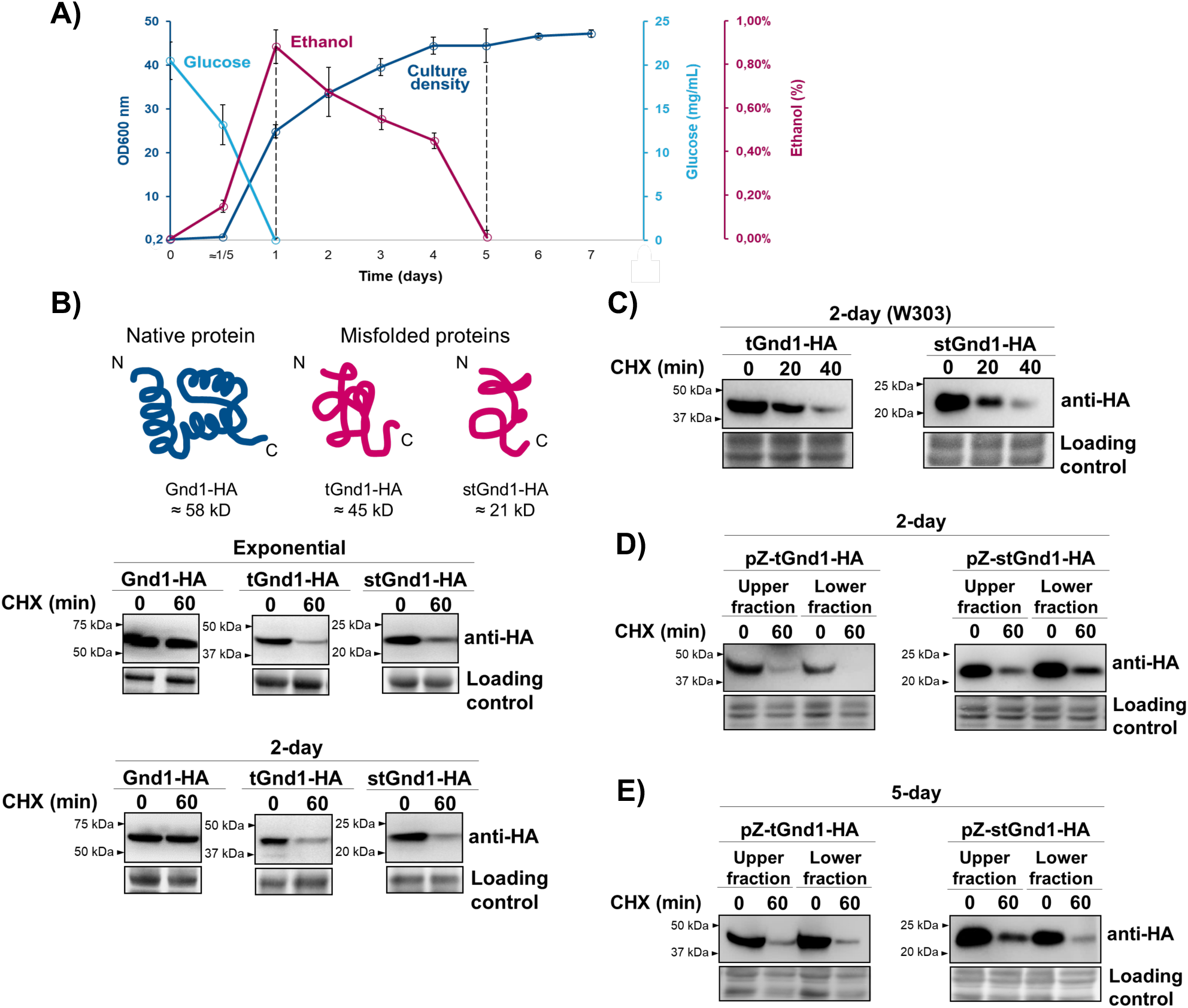
Quiescent yeast cells target misfolded proteins tGnd1 and stGnd1 for selective degradation. **A**, growth curve of yeast strain BY4741 in complete liquid medium YPD with 2 % glucose. Cells were inoculated at an initial optical density of OD_600_ 0.2 and cultured for 7 days without media change. Optical density and concentration of glucose and ethanol were measured at indicated time points. Results are presented as mean value ± standard deviation (n=3). **B-E**, protein stability was analyzed by cycloheximide (CHX) chase. Cells were collected at indicated time points after cycloheximide addition and analyzed by Western blot (anti-HA). Total proteins were visualized using stain-free technology (Bio-Rad) and used as a loading control. **B**, cells of the wild type yeast strain BY4741 expressing Gnd1-HA (DFY006), tGnd1-HA (DFY004) or stGnd-HA (DFY005) from *TEF1*-gene promoter were analyzed in exponentially growing cultures (“exponential”). Cells expressing Gnd1-HA (DFY003), tGnd1-HA (DFY001) or stGnd-HA (DFY002) from *PIR3*-gene promoter were analyzed in cultures grown for 48 hours after inoculation (“2-day”). **C**, cells of the wild type yeast strain W303 expressing tGnd1-HA (MPY166) and stGnd1-HA (MPY167) from the *PIR3*-promoter were grown for 48 hours (“2-day”) and examined as above. **D-E**, tGnd1 and stGnd1 are degraded in two-day- and five-day-old cells from both density fractions. Cells expressing tGnd1-HA (DFY052) or stGnd1-HA (DFY053) under the control of a progesterone-inducible pZ promoter were grown for two (D) or five (E) days, expression was induced by the addition of 100 μM progesterone for 60 min before performing cycloheximide chase. Cells were separated in a density-gradient and upper and lower density fractions were analyzed by Western blot.

Cultures entering quiescence are heterogeneous and two fractions can be separated based on different cell density (42). To examine whether misfolded proteins are targeted for degradation in cells from both density fractions, the cells were separated by centrifugation in the density gradient (Fig. 1D). The use of a progesterone-inducible Z-promoter enabled s/tGnd1 expression at later stages of cell quiescence. The analysis demonstrated that tGnd1 and stGnd1 undergo degradation in cells from both density fractions (Fig. 1D). Moreover, the stability of tGnd1 and stGnd1 in cells from five-day-old cultures was similar to that observed in cells from two-day-old cultures (Fig. 1E), indicating that the degradation of misfolded proteins persists throughout the later stages of quiescence.

### Protein quality control in quiescent cells involves active ubiquitin-proteasome system and spatial sequestration of misfolded proteins to inclusions

In exponentially growing cells, tGnd1 and stGnd1 are targeted for degradation by the activity of E3 ligases San1 and Ubr1 (11). To test whether selective degradation of s/tGnd1 in quiescent cells also requires the activity of the same E3 ubiquitin ligases, we examined the stability of tGnd1-HA and stGnd1-HA expressed in single *ubr1Δ* and *san1Δ* mutants, and in the double *san1Δ ubr1Δ* mutant (Fig. 2). In exponentially growing cells, stGnd1 was mainly targeted by the E3 ubiquitin ligase Ubr1, while efficient stabilization of tGnd1 required the deletion of both *SAN1* and *UBR1* (Fig. 2A, B), which is consistent with a previous study (11). In quiescent cells, in which protein expression was regulated by the *PIR3*-promoter or a progesterone-inducible Z promoter, the degradation of tGnd1 was more dependent on Ubr1, than on San1, particularly when tGnd1 was constitutively expressed from the *PIR3*-promoter (Fig. 2A). This result suggests that in quiescent cells, tGnd1 becomes less accessible to ubiquitination by San1, especially upon prolonged expression. Degradation of stGnd1 in quiescent cells remained predominantly Ubr1-dependent, as in exponentially growing cells (Fig. 2B).

**Figure 2.**
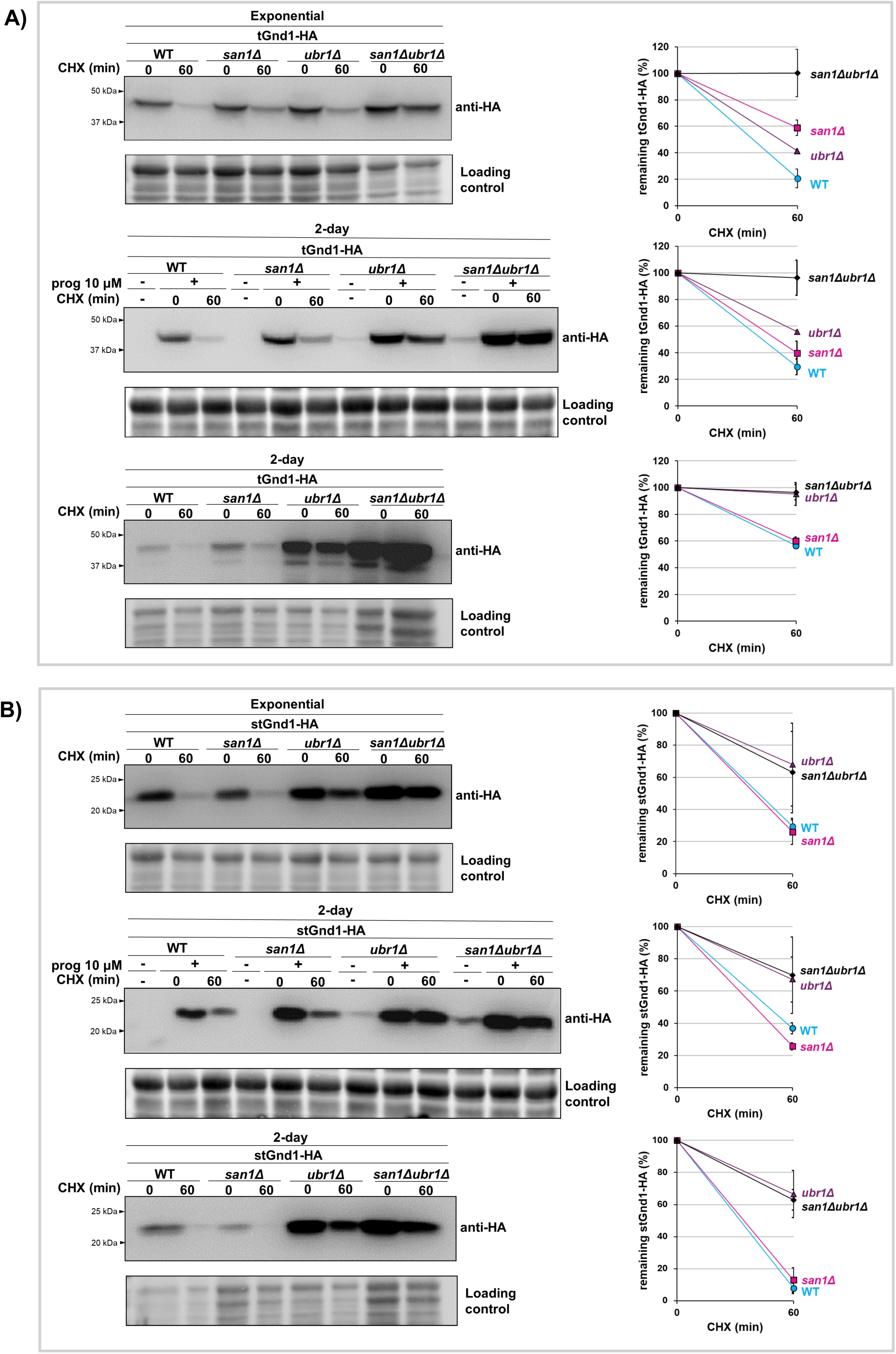
Degradation of misfolded proteins tGnd1 and stGnd1 in quiescent cells depends predominantly on Ubr1, whereas San1 plays a minor role. Western blot analysis of the cycloheximide chase (CHX). Cells from exponentially growing cultures (“exponential”) expressing indicated proteins from the constitutive *TEF1*-promoter (A-B, upper panels) and cells from 2-day old cultures expressing indicated proteins from the constitutive *PIR3*-promoter (A-B, lower panels) or progesterone-inducible pZ-promoter (A-B, middle panels) were analyzed as in Fig. 1. Graphs represent tGnd1-HA and stGnd1-HA protein levels as a percentage of the protein present at the time point 0 min. Average values and standard deviation are shown (n=2). **A, upper panel**, tGnd1-HA in the wild type BY4741 (DFY004), *san1Δ* (DFY039), *ubr1Δ* (DFY043) and *ubr1Δ san1Δ* (DFY057). **A, middle panel**, tGnd1-HA in wild-type (DFY052), *san1Δ* (DFY118), *ubr1Δ* (DFY119), and *ubr1Δ san1Δ* (DFY120). **A, lower panel**, tGnd1-HA in wild type (DFY001), *san1Δ* (DFY037), *ubr1Δ* (DFY041) and *ubr1Δsan1Δ* (DFY055). **B, upper panel**, stGnd1-HA in wild-type strain (DFY005), *san1Δ* (DFY040), *ubr1Δ* (DFY044), and *ubr1Δsan1Δ* (DFY058). **B, middle panel**, stGnd1-HA in wild-type (DFY053), *san1Δ* (DFY121), *ubr1Δ* (DFY122), and *ubr1Δ san1Δ* (DFY123). **B, lower panel**, stGnd1-HA in wild-type (DFY002), *san1Δ* (DFY038), *ubr1Δ* (DFY048), and *ubr1Δ san1Δ* (DFY056). Stain-free total protein (Bio-Rad) was used as a loading control.

A previous study has shown that tGnd1 expressed in exponentially growing cells of the *san1Δ ubr1Δ* mutant forms inclusions (22), while the localization of stGnd1 has not been previously tested. To investigate whether s/tGnd1-HA form inclusions in quiescent cells, proteins were N-terminally tagged with the green fluorescent protein ymNeonGreen (NGreen) (43). Analysis of the protein stability by Western blot demonstrated that the degradation of NGreen-tagged constructs was dependent on San1 and Ubr1 (Fig. 3A, B), in a manner similar to that previously observed for HA-tagged constructs (Fig. 2A, B).

**Figure 3.**
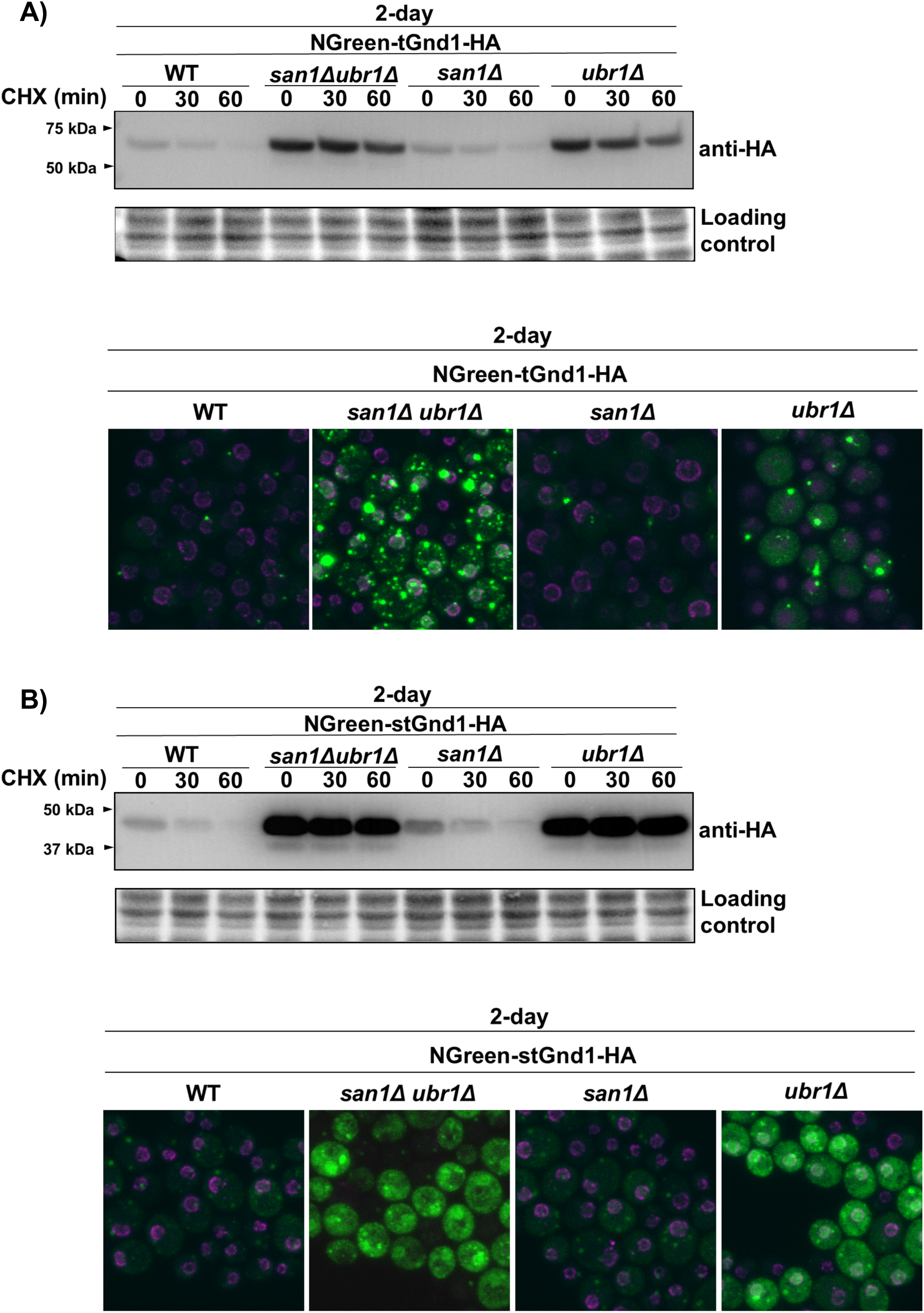
tGnd1 expressed in quiescent cells of ubiquitination mutants forms inclusions, whereas stGnd1 retains a predominantly diffuse localization. Immunoblot and localization of NGreen-tGnd1-HA (A) and NGreen-tGnd1-HA (B) in quiescent cells. NGreen-tGnd1-HA was expressed in 2-day cell cultures under the constitutive *PIR3*-promoter in wild type BY4741 (DFY192) and deletion mutants *san1Δ* (DFY193), *ubr1Δ* (DFY194), and *ubr1Δ san1Δ* (DFY195). NGreen-stGnd1-HA was similarly expressed in wild type (DFY196), *san1Δ* (DFY197), *ubr1Δ* (DFY198), and *ubr1Δ san1Δ* (DFY200). Cycloheximide chase analysis was performed as described in Fig. 1. Stain-free total protein (Bio-Rad) was used as a loading control. Protein localization of the same cultures as in A was analyzed by confocal fluorescent microscopy. The green signal represents green fluorescent protein ymNeonGreen (NGreen), while red signal represents nucleoporin Nup49-mScarlet. Shown is the whole z-stack.

The fluorescent signal of NGreen-tGnd1 and NGreen-stGnd1 expressed in the quiescent cells of the wild-type strain was weak. Nevertheless, a small percentage of the cells expressing NGreen-tGnd1, exhibited discrete puncta (Fig. 3A), while NGreen-stGnd1 localized diffusely (Fig. 3B). In the *san1Δ* mutant, the localization of tGnd1 was similar to that observed in the wild-type strain, consistent with the lack of protein stabilization in the single *san1Δ* mutant (Fig. 3A, upper panel). In contrast, in the *ubr1Δ* mutant, tGnd1 formed large inclusions, and their number and the size were further increased in the double *san1Δ ubr1Δ* mutant (Fig. 3A). Furthermore, in addition to a single large inclusion of tGnd1, several smaller puncta were clearly visible in the *san1Δ ubr1Δ* mutant cells. The appearance of the small puncta is consistent with the ongoing protein synthesis of NGreen-tGnd1-HA in our experimental setup, and with the previous reports of Q-bodies, small dynamic structures that initially sequester misfolded proteins and eventually coalesce into larger inclusions (23, 24).

In contrast to NGreen-tGnd1, which formed distinct inclusions, NGreen-stGnd1 retained a predominantly diffuse localization, even in the ubiquitination mutants *ubr1Δ* and *san1Δ ubr1Δ*, which exhibit protein stabilization on Western blot (Fig. 3B). In accordance with the dependence of stGnd1 degradation on Ubr1, the double mutation *san1Δ ubr1Δ* did not result in an additive effect. In addition to its predominantly diffuse localization, small granules of stGnd1 could also be detected, however, their low intensity and appearance was clearly different from the large and prominent inclusions of tGnd1. In summary, tGnd1 localizes to the inclusions, which exhibit a considerable increase in size and frequency in the ubiquitination mutants, whereas stGnd1 localization remains predominantly diffuse even in mutants with impaired protein degradation.

Next, we set out to test whether degradation of misfolded proteins in quiescent cells involves 26S proteasomes. In contrast to the dividing yeast cells, in which the majority of the proteasomes localize to the nucleus (36), in quiescent cells a large pool of the proteasomes relocalizes to the nuclear periphery and to the cytoplasmic storage granules, which are thought to contain inactive proteasomes that are disassembled into core and regulatory particles (38, 39). To investigate whether quiescent cells retain proteasomal degradation of misfolded proteins, we examined the degradation of s/tGnd1 in the proteasomal mutant *rpn11-m1* (44), which expresses non-functional Rpn11, a deubiquitinase that is critical for the functioning of the 26S proteasome (20) (Fig. 4). The *rpn11-m1* mutant is temperature sensitive, therefore the cells were grown at 25°C, and shifted to 37°C for 30 minutes prior to the cycloheximide chase. The analysis demonstrated that tGnd1 and stGnd1 were stabilized in the *rpn11-m1* mutant, even in cells incubated at 25°C (Fig. 4A, B), indicating the crucial role of fully assembled and functionally active 26S proteasome in the degradation of misfolded proteins in quiescent cells. Together the data indicate that the 26S proteasomes remain actively engaged in the degradation of quality control substrates during cell quiescence.

**Figure 4.**
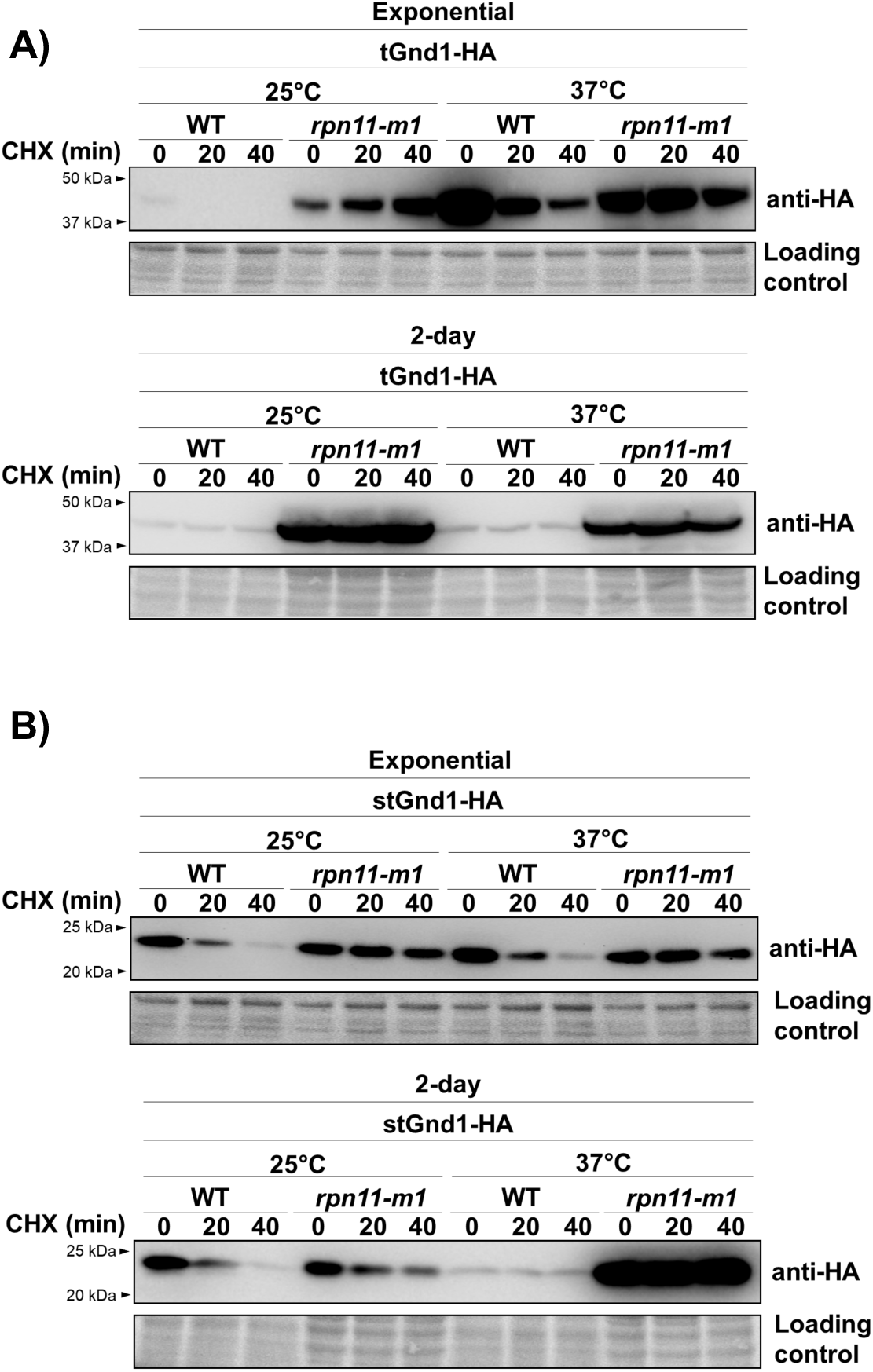
The degradation of misfolded proteins tGnd1 and stGnd1 in quiescent cells is dependent on the activity of the proteasome. Stability of tGnd1-HA and stGnd1-HA was assessed in the temperature-sensitive proteasome mutant strain *rpn11-m1*. Cell cultures were grown at permissive temperature of 25°C to the exponential phase or for 2 days, then shifted to a restrictive temperature of 37°C for 30 min, followed by cycloheximide chase and Western blot analysis as described in Fig. 1. tGnd1-HA (A) and stGnd1-HA (B) were expressed from centromeric plasmids under the control of *TEF1*-promoter (pMB214, pMB215) or *PIR3-* promoter (pMB211, pMB212) in the wild type (W303) and *rpn11-m1* mutant (YP337). Stain-free total protein (Bio-Rad) was used as a loading control.

### Degradation of certain misfolded proteins in quiescent cells requires Cue5-independent autophagy pathway

Cells cultured in a rich glucose-based medium gradually exhaust glucose and switch to using accumulated ethanol as a carbon source (31). Autophagy is induced starting at the ethanol-utilizing phase (45), and can be observed through the accumulation of free GFP in cells expressing GFP-Atg8 (46) (Fig. 5A). To examine whether the degradation of misfolded proteins in quiescent yeast cells requires functional autophagy, we compared the stability of tGnd-HA and stGnd1-HA expressed in wild-type and autophagy-deficient *atg1Δ* and *atg8Δ* mutant strains. The lack of Atg1 and Atg8 did not affect the degradation of stGnd1, indicating that autophagy is not a major pathway for the degradation of stGnd1 in quiescent cells (Fig. 5C). Stability of a cytoplasmic enzyme Pgk1 was also unaffected. In contrast, tGnd1 was stabilized in both *atg1Δ* and *atg8Δ* mutant strains (Fig. 5B), indicating the critical requirement for the functional autophagy in the degradation of tGnd1. A similar stabilization of tGnd1 was observed in the *atg1Δ* mutant of another strain background, W303 (Fig. S2).

**Figure 5.**
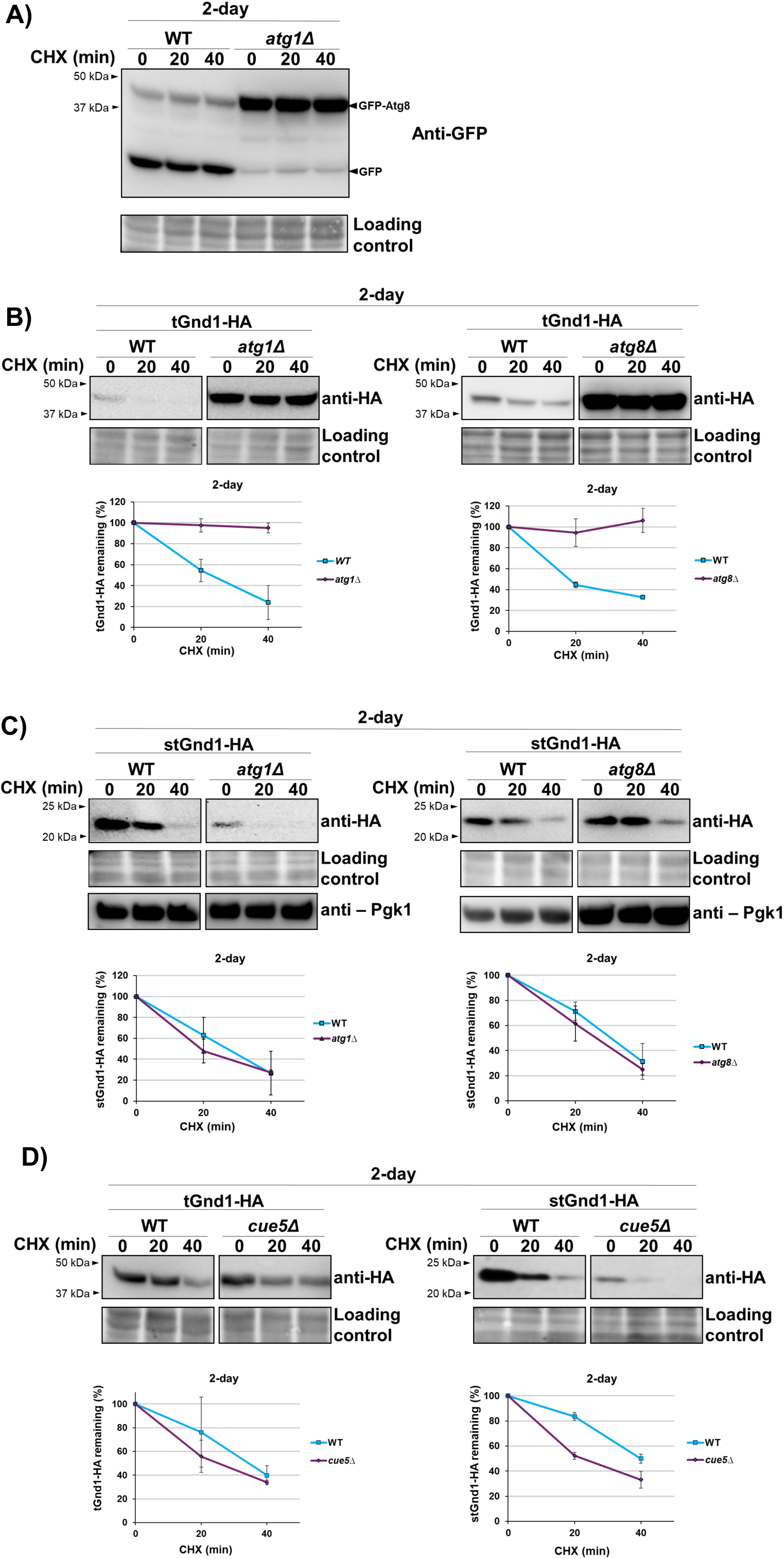
The degradation of misfolded protein tGnd1, but not stGnd1, is impaired in the quiescent cells of autophagy mutants *atg1Δ* and *atg8Δ*. Western blot analysis of the cycloheximide chase was performed as in Fig.1. **A**, accumulation of free GFP in the wild-type (MBY501) and *atg1Δ* mutant (MBY507) strains expressing GFP-Atg8 was analyzed by Western blot (anti-GFP). **B-D** degradation of tGnd1-HA and stGnd1-HA expressed from *PIR3*-promoter was analyzed in the wild-type (MBY513, MBY514), *atg1Δ* (MBY483, MBY487) and *atg8Δ* (MBY484, MBY488) strains, and in the *cue5Δ* mutant (MBY482, MBY486). Stain-free total protein (Bio-Rad) was used as a loading control. The samples shown were present on the same membrane and imaged simultaneously. Graphs represent tGnd1-HA and stGnd1-HA protein levels as a percentage of the protein present at the time point 0 min. Average values and standard deviation are shown (n=2).

The data showing that tGnd1, but not stGnd1 or Pgk1, is stabilized in autophagy mutants, suggested degradation selectivity. In yeast, selective autophagy of aggregation-prone proteins is mediated by the ubiquitin- and Atg8-binding protein Cue5, the only known ubiquitin-binding autophagy receptor in this organism (47, 48). To test the involvement of Cue5, we examined the stability of tGnd1 in a *cue5Δ* deletion mutant, however there was no effect (Fig. 5D), suggesting a distinct, Cue5-independent, mechanism for autophagy selectivity.

### The clearance of specific misfolded proteins in quiescent cells critically depends on the intact nucleus-vacuole junctions

A recent study showed that the clearance of deposition sites INQ and JUNQ involves their vacuolar targeting through the nucleus-vacuole junction (NVJ) (24), a site formed by direct interactions between the vacuolar membrane protein Vac8 and the outer nuclear membrane protein Nvj1, whose size and frequency increases upon starvation (49, 50). To test the possibility that the clearance of misfolded proteins in quiescent cells involves nucleus-vacuole junctions, we examined the stability of tGnd1 and stGnd1 in the *nvj1Δ* and *vac8Δ* mutants, which are unable to form NVJ (49). The degradation of tGnd1 was clearly impaired in the quiescent cells of the *nvj1Δ* and *vac8Δ* mutants, indicating a critical role of the NVJ in the clearance of tGnd1 (Fig. 6A, lower left panel). In contrast, the degradation of stGnd1 was not affected by the NVJ mutants (Fig. 6A, lower right panel). Considering the localization of tGnd1, but not stGnd1, to the inclusions, our data are consistent with a previous study (24) showing the localization of INQ/ JUNQ in the vicinity of the NVJ and the delivery of the INQ/JUNQ-localized misfolded proteins to the vacuole. Importantly, in our experiments, neither tGnd1 nor stGnd1 exhibited a detectable stabilization in the exponentially growing cells of the *nvj1Δ* and *vac8Δ* mutants (Fig. 6A, upper panels), suggesting that the role of NVJ in the clearance of misfolded proteins becomes prominent upon cell entry into quiescence.

**Figure 6.**
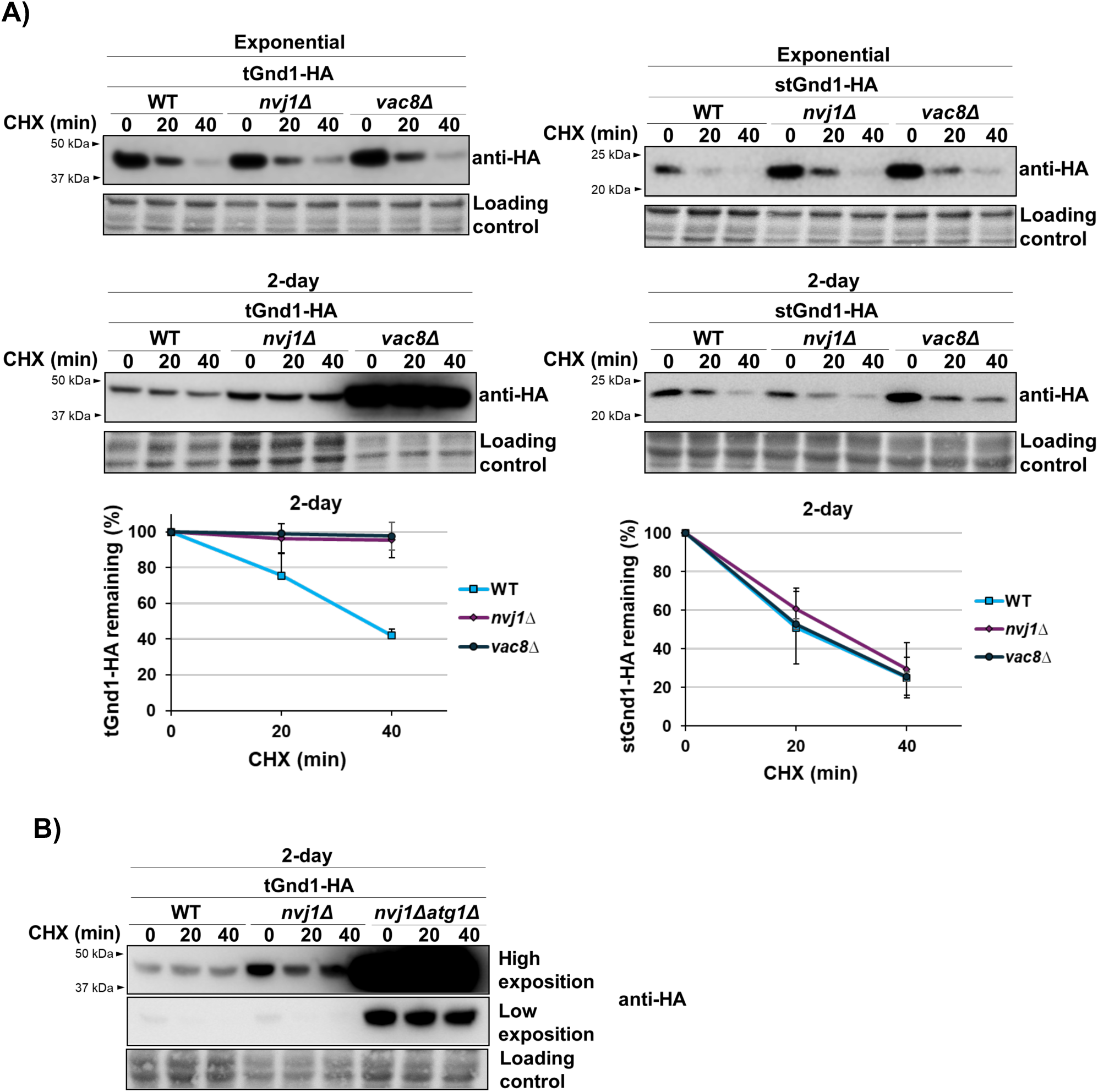
Misfolded protein tGnd1 is stabilized in the nucleus-vacuole junction mutant *nvj1Δ*, and double mutation *nvj1Δ atg1Δ* leads to an additive effect. Western blot analysis of the cycloheximide chase. **A**, Cells expressing tGnd1-HA or stGnd1-HA under the control of *TEF1*-promoter (exponentially growing culture) or *PIR3*-promoter (2-day old culture) in the wild type (MPY152, MPY153, MPY154, MPY155), *nvj1Δ* mutant (MPY156, MPY158, MPY160, MPY162), and *vac8Δ* mutant (MPY157, MPY159, MPY161, MPY163) were analyzed as in Figure 1. **B**, the stability of tGnd1-HA was compared in the 2 days-old cultures from single *nvj1Δ* (MPY156) and double *nvj1Δ atg1Δ* (MPY164) mutant. Stain-free total protein (Bio-Rad) was used as a loading control. Graphs represent tGnd1-HA and stGnd1-HA protein levels as a percentage of the protein present at the time point 0 min. Average values and standard deviation are shown (n=2).

The stabilization of tGnd1 in the quiescent cells of the *vac8Δ* mutant was significantly stronger than in the *nvj1Δ* mutant. This data indicated that the loss of Vac8 leads to the impairment of an additional pathway that is involved in tGnd1 degradation. Vac8 has been shown to be required for both bulk and selective autophagy (51–53). To test whether the clearance of tGnd1 via NVJ-dependent degradation and autophagy represent two separate pathways, we compared tGnd1 degradation in the single *nvj1Δ* mutant and a double *nvj1Δ atgj1Δ* mutant and examined whether the double *nvj1Δ atgj1Δ* mutant results in an additive effect. The stabilization of tGnd1 was considerably higher in the *nvj1Δ atgj1Δ* double mutant, than in the single *nvj1Δ* mutant (Fig. 6B), demonstrating an additive effect of impaired autophagy and disrupted NVJ. The data indicate that the NVJ-dependent clearance and autophagy represent two separate pathways in the degradation of tGnd1, and suggest that the two pathways operate in parallel.

## Discussion

This study shows that quiescent yeast cells retain degradation-mediated protein quality control, employing a combination of different pathways to mitigate the accumulation of misfolded proteins. Despite the previously reported relocalization of proteasomes in quiescent cells, our data indicate that a significant pool of fully assembled and active 26S proteasomes are engaged in the degradation of quality control substrates. Moreover, in contrast to the exponentially growing cells, the efficient clearance of certain substrates necessitates the presence of intact nucleus-vacuole junctions and autophagy, which is independent of the only known ubiquitin-binding autophagy receptor in yeast, Cue5.

Previous reports of the proteasomal substrates of the (that are ubiquitnated by the) N-end rule (54) and ubiquitin fusion degradation (UFD) pathway (55) pathways that were expressed in cells from stationary phase cultures suggested that the substrate degradation in quiescent cells was greatly decreased (41) or even abolished (39). However, due to the hardly detectable expression of the model substrates under the examined conditions (39), the interpretation was difficult. Furthermore, an earlier study of cells from stationary phase culture showed that a thermosensitive luciferase mutant formed inclusions, which suggested that quiescent cells may manage misfolded proteins primarily by sequestration. However, the role of degradation-mediated PQC in this process was not investigated. In our study, we expressed model misfolded proteins from the constitutive and inducible promoters that are active in quiescent cells and showed that misfolded proteins are targeted for selective degradation, in both early and later phases of cell quiescence.

A recent study has reported that the yeast strain BY4741 is unable to enter quiescence from the rich medium (56). In that study, BY4741 ceased dividing several hours before glucose exhaustion and the cell density reached a plateau at a low cell density (OD_600_ < 10), already at 20 hours post-inoculation. This was in contrast to the yeast strain W303, which did not exhibit such phenotype (56). In our experiments, BY4741 continued to grow after glucose exhaustion, reaching a high cell density by the fourth day post-inoculation (OD_600_ above 40). Therefore, the growth curve for BY4741 observed in our study is similar to that reported for W303 by the Breeden laboratory. The discrepancy in the growth of BY4741-derived strains between the Breeden laboratory and our laboratory may be attributed to the differences in the composition of the growth media. Nevertheless, we have performed the key experiments in the W303 strain background. The results demonstrated that misfolded proteins s/tGnd1 were degraded in W303 in a proteasome-dependent manner, and that tGnd1 was stabilized in an autophagy mutant. These findings indicated that active degradation pathways are present in the W303 strain, in a manner similar to those observed in BY4741-derived strains.

In quiescent cells, proteasomes relocalize to the nuclear periphery and to cytoplasmic storage granules (37), which are thought to contain disassembled, inactive, proteasomes (38, 39). Here we showed that the efficient removal of the misfolded protein stGnd1 in quiescent cells was Rpn11-dependent, and did not require other pathways, such as autophagy or NVJ-mediated clearance. This finding supports the presence of fully assembled and functionally active 26S proteasomes in quiescent cells, and is consistent with a previous report showing that assembled 26S proteasomes were still detectable in quiescent cells (37). Our results indicate that 26S proteasomes are actively involved in the degradation of misfolded proteins in quiescent cells, suggesting that the pool of the proteasomes that is not integrated into proteasome storage granules is sufficient to meet the demand for PQC during quiescence, a state characterized by downregulation of new protein synthesis, and thus a lower burden of misfolded proteins.

Our results demonstrated that tGnd1, but not stGnd1, was stabilized in quiescent cells of autophagy mutants, indicating selectivity. Inactivation of the only known ubiquitin-binding autophagy receptor in yeast, Cue5, had no effect on tGnd1 stability, suggesting that a distinct mechanism is involved. The sequestration of tGnd1, but not stGnd1, into the inclusions suggests that autophagy selectivity may be contingent upon tGnd1 localization to the inclusions. A recent report has showed Cue5-independent autophagy of a poly-glutamine expanded huntingtin (HttQ103) protein expressed in dividing yeast cells (57). The process was termed inclusion body autophagy or IBophagy, and was dependent on the selective autophagy receptors Atg36, Atg39 and Atg40 (57). It is possible that a similar process is involved in degradation of tGnd1 in quiescent cells.

In our experiments tGnd1 was clearly stabilized in the nucleus-vacuole junctions (NVJ) mutants, but only in quiescent cells. This finding is also consistent with previous studies that have demonstrated an increase in the frequency and size of the NVJ increases upon prolonged growth in YPD (50). The lack of tGnd1 stabilization in the NVJ mutants of the exponentially growing cells suggests that the role of the NVJ-mediated clearance becomes critical in cell quiescence, presumably due to the limits in the proteasomal degradation of certain substrates.

Notably, tGnd1 exhibited stabilization in the autophagy- and NVJ-mutants that had a functional ubiquitin-proteasome system, suggesting that proteasomes, autophagy and NVJ-dependent clearance, function in parallel. The finding that the clearance of tGnd1 in quiescent cells additionally depends on autophagy and NVJ-dependent clearance suggests that the efficiency of proteasomal degradation of tGnd1 in quiescent cells is limited. The observation that another substrate, stGnd1, did not require NVJ or autophagy for efficient clearance, strongly suggests that the proteasomes themselves are not the limiting factor. Rather, substrate-specific factors that are necessary to ensure the delivery of the protein to the proteasome may be affected.

Our results suggest that quiescent cells utilize similar Ubr1- and San1-dependent ubiquitination pathways for tagging misfolded proteins for degradation as exponentially growing cells. The observed activity of Ubr1 in quiescent cells is also consistent with an earlier observation of increased steady-state levels of a model misfolded protein ΔssCL*myc in *ubr1Δ* mutant cells from stationary phase cultures (58). Of note, in quiescent cells, the degradation of tGnd1 was more dependent on Ubr1 than on San1. This is in contrast to exponentially growing cells in which San1 had a similar contribution as Ubr1. This finding suggests that in quiescent cells, the delivery of tGnd1 to San1 for ubiquitination may be impaired, potentially due to an inability to transport tGnd1 into the nucleus, or a lack of chaperones.

In support of the possibility that degradation of tGnd1 in quiescent cells is limited by the availability of chaperones, the efficient degradation of tGnd1 in exponentially growing cells has been shown to require Hsp40 chaperone Sis1, whereas stGnd1 degradation was largely Sis1-independent (59). Sis1 facilitates the delivery of misfolded proteins into the nucleus (59, 60) and nuclear substrate ubiquitination (15). Sis1 has also been implicated in Ubr1-mediated degradation (59), although its role in this process appears less prominent than its role in the nuclear import. Additionally, a recent report demonstrated that the transition to respiratory metabolism and the accompanying decrease in translation rates in quiescent cells leads to the re-localization of protein disaggregase Hsp104 to the nucleus (61). It is plausible that the reduced levels of Hsp104 in the cytoplasm result in a lack of Hsp104 availability for the disaggregation of proteasomal substrates in the cytoplasm.

The data on PQC pathways in quiescent cells of mammalian organisms is limited. A transcriptomic study of mouse neural stem cells has shown that activated neural stem cells exhibit increased expression of proteasome-associated genes, while quiescent neural stem cells exhibit increased expression of lysosome-associated genes, and have fewer catalytically active proteasomes (62). Quiescent neural stem cells accumulated insoluble protein aggregates, however protein degradation has not been tested. In line with that, a study investigating dermal fibroblasts showed that dermal fibroblasts increase degradation rate of long-lived proteins upon entry into quiescence, by activating lysosome biogenesis and upregulating autophagy (63). Thus, autophagy upregulation upon cell cycle exit may be a common feature of quiescent cells in mammalian organisms. Collectively, the available data from mammalian organisms are in line with our results in yeast, which indicate that in addition to misfolded protein degradation via ubiquitin-proteasome system, quiescent cells need functional autophagy to maintain protein homeostasis. Together, our findings underscore the importance of misfolded protein clearance during cell quiescence, and contribute to understanding the interplay of different degradation pathways.

## Experimental procedures

### Yeast strains

Yeast *Saccharomyces cerevisiae* strain used in this study are listed in Table 1. All strains used in this study were isogenic to BY4741 (64), except MPY166, *atg1Δ* mutant strain MPY170 and *rpn11-m1* mutant strain YP337, which were derived from W303. The description of strain construction is provided in Table 2. Strains were constructed by homologous recombination of DNA constructs or plasmids cut with the indicated restriction enzyme that were transformed into yeast strains. Molecular cloning was performed by standard methods. Sequences of all primers used in this study are listed in the Table 3. Genome integration of the transformed DNA constructs was verified by PCR.

### Yeast culture and growth media

Standard yeast culture media, such as ammonia-based synthetic complex dextrose (SC) medium containing 2% D-glucose were used. Yeast extract, peptone, dextrose (YPD) medium was prepared from 1 % yeast extract (YEA03, Formedium Ltd. England), 2 % peptone (PEP03, Formedium Ltd. England) and 2 % D-glucose (GLU03, Formedium Ltd. England). Antibiotic selections were made on solid YPD containing 200 mg/ l G418 or 300 mg/l hygromycin B. When grown in liquid media, culture tubes and baffled flasks with loose-fitting caps were used, and cells were incubated in an orbital shaker (Innova 40R, New Brunswick) with shaking at 240 rpm. Cells were grown at 30°C unless indicated otherwise. For the analysis of exponentially growing and quiescent cells, yeast overnight cultures were diluted in fresh YPD containing 2 % glucose to an optical density OD_600_ of 0.2 and grown to the exponential phase (OD_600_ 0.8-1.0), or for a period of two to seven days, as indicated, without media change. Where indicated, Z promoter was activated by the addition of progesterone (Sigma-Aldrich, St. Louis, Missouri, USA) to the media at the final concentration of 10 μM or 100 μM, as indicated, for one hour.

### Plasmids

All plasmids used in this study are listed in Table 4. The description of the plasmid construction is listed in Table 5. Sequences of primers used for construction are listed in Table 3.

### Measurement of glucose and ethanol concentration

Glucose concentration in the medium was determined using the Glucose (GO) Assay Kit (Sigma-Aldrich, St. Louis, Missouri, USA) following the manufacturer’s instructions. Ethanol concentration was measured using a previously described protocol (65). Values of measurements of three independent samples are presented as mean values with standard deviations.

### Cell fractionation by centrifugation in density gradient

Cell fractionation in Percoll (Cytiva, Washington DC, USA) density gradient was performed as previously described (42). Gradient was prepared by mixing Percoll with 1.5 M NaCl in a 9:1 volume ratio, resulting in a total volume of 10 mL and a final NaCl concentration of 167 mM. To form the gradients, 1.8 mL of the Percoll solution was placed into 2 mL tubes and centrifuged (12000 RPM, 15 min, 4°C). Yeast overnight cultures were diluted in fresh YPD containing 2 % glucose to an optical density OD_600_ of 0.2 and grown for two or five days. A cell culture volume corresponding to 36 OD_600_ units of cells was harvested by centrifugation (3000 x g, 3 min), resuspended in 180 μl 50 mM Tris buffer, and overlaid onto the preformed Percoll density gradient in 2 ml tubes. Cells were centrifuged in Percoll density gradient (400 x g for 60 min at 20°C). Cell fractions were collected by pipetting, washed in 8 mL of Tris-buffer, and resuspended in 1 mL Tris-buffer. The optical density (OD_600_) of each cell fraction was measured, and a volume corresponding to 1 OD_600_ units of cells was pelleted. Protein extraction and analysis by Western blot was performed as described below.

### Cycloheximide chase and Western blot analyses

Yeast overnight cultures were diluted in fresh YPD containing 2 % glucose to an optical density OD_600_ of 0.2 and grown to the exponential phase (OD_600_ 0.8-1.0), or for a period of two to five days, as indicated. Translation inhibitor cycloheximide (Sigma-Aldrich, St. Louis, Missouri, USA) was added to the cell culture at a final concentration of 100 µg/mL. Cells were collected at the indicated time points following cycloheximide addition.

Total cell lysates were prepared as described (66) with some modifications. For the analysis of the proteins from exponentially growing cultures (OD_600_ 0.8-1.0), 1.5 mL of cell culture was harvested. Cells were collected by centrifugation for (5 min, 11000 RPM, 4 °C), the cell pellet was frozen in liquid nitrogen and stored at −20 °C until samples from all time points were collected. Protein isolation was done based on a previously described protocol (2) with some modifications. Briefly, cells were resuspended in 100 µL of ice-cold water, 100 µL of ice-cold 0.2M NaOH was added, followed by incubation on ice for 10 min and centrifugation (5 min, 11000 RPM at 4 °C). Cell pellets were resuspended in 50 µL SDS buffer (0.06 M Tris±HCl, pH 6.8, 5 % glycerol, 2 % SDS, 4 % β-mercaptoethanol, 0.0025 % bromophenol blue), and incubated at 95 °C for 3 min. For the analysis of the proteins from density fractions isolated as above, the pellet was resuspended in 20 uL of SDS-buffer. Samples were centrifuged (5 min, 11000 RPM at 23 °C) and the supernatant was kept. For the analysis of the proteins from quiescent cells (two- and five-day-old cultures), a cell culture volume corresponding to 10 OD_600_ units of cells was harvested, and the same protocol as above was applied, but using double volumes of reagents, due to a larger number of cells (200 µL of distilled water, 200 µL of 0.2 NaOH, and 100 µL of SDS buffer).

Western blot was performed by antibodies: anti-HA (12CA5, Ogris laboratory, Max F. Perutz Laboratories, Vienna, Austria, 1/1000), anti-Pgk1 (22C5, RRID: AB_2546088, Invitrogen 1/20,000), anti-GFP (Roche Diagnostics GmbH, Germany, 1/1000), and HRP-conjugated secondary antibodies Cell Signaling Technology, Massachusetts, USA, (#7076, 1/2000). Chemiluminescent signal intensity of immunoreactive bands was imaged by ChemiDoc MP Imaging System (BioRad Laboratories) and quantified by ImageLab (BioRad Laboratories). Stain-free signal of the total proteins (BioRad Laboratories) was used as a loading control. A representative band of the stain-free signal is shown in figures. Signal intensities of anti-HA-immunoreactive bands representing tGnd1-HA and stGnd1-HA were normalized to the Stain-free signal (BioRad Laboratories) of the total proteins originating from the whole lane. At least two independent samples were analyzed. Signal intensities are presented as mean values with standard deviations.

### Microscopy

Yeast overnight cultures were diluted in fresh YPD containing 2 % glucose to an optical density OD_600_ of 0.2 and grown for two days. Cells were fixed in 0.8 % formaldehyde for 10 min, centrifuged, and washed twice in phosphate-buffered saline (PBS). Cell pellet was resuspended in PBS and dropped onto coverslips pre-coated with concanavalin-A (Sigma-Aldrich). Images were captured using confocal fluorescent microscope Olympus FV3000 (Olympus, Tokio, Japan) fitted with an Olympus-DP74 digital camera, an Olympus 60× oil-immersion objective (Olympus UPlanSApo 60x/1.35 Oil Microscope Objective), and FV31S-SW Fluoview program. Figures were prepared using Fiji (ImageJ). The whole z-stack of the cells is shown.

## Data availability

All data are contained within the manuscript.

## Supporting information

This article contains supporting information (Figure S1, Figure S2).

## Acknowledgments

Strains yTB281 and yTB293 were a kind gift from Claudine Kraft. Strains W303 and YP337 were a kind gift from Elah Pick. Plasmids pHES830 and pHES836 were a gift from Hana El-Samad, plasmids pRG206MX and pRG216 were a gift from Joerg Stelling, plasmid pXP732 was gift from Nancy Da Silva, and pFA6a-mScarlet was a gift from Bas Teusink, all via Addgene.

