## Supplemental Figures for "Quiescent cells maintain active degradation-mediated protein quality control requiring proteasome, autophagy and nucleus-vacuole junctions"

### Supporting information

**Figure S1.**

**Figure S2.**

### Supplementary Figure S1

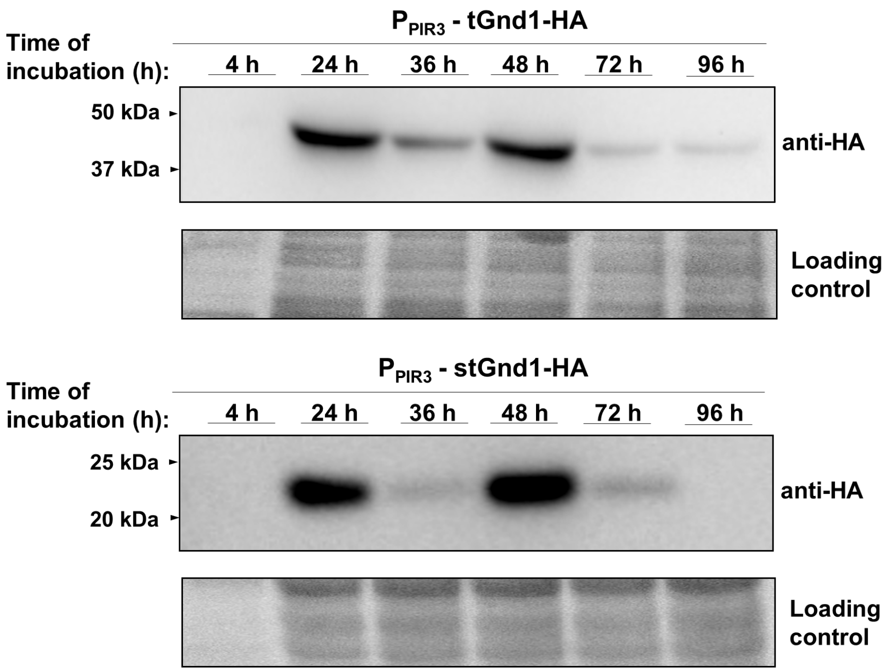


**Figure S1. Expression of model misfolded proteins tGnd1-HA and stGnd1-HA under the control of constitutive *PIR3*-gene promoter in proliferating and quiescent cells.** Western blot analysis of tGnd1-HA (DFY001) and stGnd1 (DFY002) expressed under the control of *PIR3*- promoter in the wild type BY4741 strain. Cells were inoculated at an initial optical density ($OD_{600}$) of 0.2 and cultured for an indicated time period. Cell lysates were analyzed by Western blot (anti-HA). Stain-free total protein (Bio-Rad) was used as a loading control.

**Supplementary Figure S2**

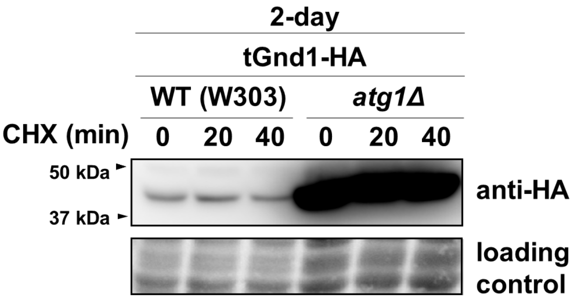


**Figure S2. Misfolded protein tGnd1-HA is stabilized in quiescent cells of the *atg1Δ* mutant of the W303 strain background.** Western blot analysis of cycloheximide chase (performed as in Fig.1). The stability of tGnd1-HA in wild-type (MPY166) and *atg1Δ* mutant (MPY170) cells from 2 days old cultures was analyzed. Stain-free total protein (Bio-Rad) was used as a loading control.
