## Supplemental Tables for "Quiescent cells maintain active degradation-mediated protein quality control requiring proteasome, autophagy and nucleus-vacuole junctions"

### Table 1. Yeast strains used in this study

| Yeast strain | Genotype | Reference |
|---|---|---|
| **W303** | *MATa; leu2-3,112; trp1-1; can1-100; ura3-1; ade2-1; his3-11,15; phi+* | (67) |
| **YP337** | *MAT a; his3_-200; ade2-101; leu2_1; ura3-52; lys2-801; trp1-62; YFR004W::rpn11-m1* | (67) |
| **BY4741** | *MATα; his3Δ1; leu2Δ0; met15Δ0; ura3Δ0* | Euroscarf (Germany) |
| **Y00253** | *his3Δ1; leu2Δ0; met15Δ0; ura3Δ0; vac8Δ::kanMX4* | Euroscarf (Germany) |
| **Y01818** | *his3Δ1; leu2Δ0; met15Δ0; ura3Δ0; cue5Δ::kanMX4* | Euroscarf (Germany) |
| **Y04547** | *his3Δ1; leu2Δ0; met15Δ0; ura3Δ0; atg1Δ::kanMX4* | Euroscarf (Germany) |
| **Y03104** | *his3Δ1; leu2Δ0; met15Δ0; ura3Δ0; atg8Δ::kanMX4* | Euroscarf (Germany) |
| **Y02889** | *his3Δ1; leu2Δ0; met15Δ0; ura3Δ0; nvj1Δ::kanMX4* | Euroscarf (Germany) |
| **Y04077** | *ura3Δ0; leu2Δ0; his3Δ1; met15Δ0; san1Δ::kanMX* | Euroscarf (Germany) |
| **Y04814** | *ura3Δ0; leu2Δ0; his3Δ1; met15Δ0; ubr1Δ::kanMX* | Euroscarf (Germany) |
| **yTB281** | *his3D1; leu2D0; ura3D0; met15D0; GFP-ATG8* | (68) |
| **yTB293** | *his3D1; leu2D0; ura3D0; met15D0; atg1::KAN; GFP-ATG8* | (Torggler et al., 2016) |
| **DFY001** | *MATα; his3Δ1; leu2Δ0; met15Δ0; ura3Δ0::PPIR- tGnd1-HA URA3* | This study |
| **DFY002** | *MATα; his3Δ1; leu2Δ0; met15Δ0; ura3Δ0::PPIR3-stGnd1-HA URA3* | This study |
| **DFY003** | *MATα; his3Δ1; leu2Δ0; met15Δ0; ura3Δ0::PPIR- Gnd1-HA URA3* | This study |
| **DFY004** | *MATα; his3Δ1; leu2Δ0; met15Δ0; ura3Δ0::PTEF1-tGnd1-HA URA3* | This study |
| **DFY005** | *MATa his3Δ1 leu2Δ0 met15Δ0 ura3Δ0::PTEF1-stGnd1-HA URA3* | This study |
| **DFY006** | *MATa his3Δ1 leu2Δ0 met15Δ0 ura3Δ0::PTEF1-Gnd1-HA URA3* | This study |
| **DFY037** | *MATa; his3Δ1; leu2Δ0; met15Δ0; ura3Δ0::PPIR3- tGnd1-HA, URA3; san1Δ::kanMX4* | This study |
| **DFY038** | *MATa; his3Δ1; leu2Δ0; met15Δ0; ura3Δ0::PPIR3-stGnd1-HA, URA3; san1Δ::kanMX4* | This study |
| **DFY039** | *MATa; his3Δ1; leu2Δ0; met15Δ0; ura3Δ0::PTEF1-tGnd1-HA, URA3; san1Δ::kanMX4* | This study |
| **DFY040** | *MATa; his3Δ1; leu2Δ0; met15Δ0; ura3Δ0::PTEF1-stGnd1-HA, URA3; san1Δ::kanMX4* | This study |
| **DFY041** | *MATa; his3Δ1; leu2Δ0; met15Δ0; ura3Δ0:: PPIR3-tGnd1-HA, URA3; ubr1Δ::kanMX4* | This study |
| **DFY042** | *MATa; his3Δ1; leu2Δ0; met15Δ0; ura3Δ0:: PPIR3-stGnd1-HA, URA3; ubr1Δ::kanMX4* | This study |
| **DFY043** | *MATa; his3Δ1; leu2Δ0; met15Δ0; ura3Δ0:: PTEF1-tGnd1-HA, URA3; ubr1Δ::kanMX4* | This study |
| **DFY044** | *MATa; his3Δ1; leu2Δ0; met15Δ0; ura3Δ0:: PTEF1-stGnd1-HA, URA3; ubr1Δ::kanMX4* | This study |
| **DFY049** | *MATa; his3Δ1; leu2Δ0; met15Δ0; ura3Δ0:: PPIR3-prog-inducible synTA-Zif268 DBD* | This study |
| **DFY050** | *MATa; his3Δ1; leu2Δ0; met15Δ0; ura3Δ0; san1Δ::kanMX4 ; ubr1Δ::hphMX* | This study |
| **DFY052** | *MATa his3Δ1:: pZ-tGnd1-HA, leu2Δ0 met15Δ0 ura3Δ0:: PPIR3-prog-inducible synTA-Zif268 DBD* | This study |
| **DFY053** | *MATa his3Δ1:: pZ-stGnd1-HA, leu2Δ0 met15Δ0 ura3Δ0:: PPIR3-prog-inducible synTA-Zif268 DBD* | This study |
| **DFY054** | *MATa his3Δ1:: pZ-Gnd1-3HA, leu2Δ0 met15Δ0 ura3Δ0:: PPIR3-prog-inducible synTA-Zif268 DBD* | This study |

| | | |
|---|---|---|
| **DFY055** | *MATa; his3Δ1; leu2Δ0; met15Δ0; ura3Δ0:: PPIR3-tGnd1-HA; san1Δ::kanMX4 ; ubr1Δ::hphMX* | This study |
| **DFY056** | *MATa; his3Δ1; leu2Δ0; met15Δ0; ura3Δ0:: PPIR3-stGnd1-HA; san1Δ::kanMX4 ;ubr1Δ::hphMX* | This study |
| **DFY057** | *MATa; his3Δ1; leu2Δ0; met15Δ0; ura3Δ0:: PTEF1-tGnd1-HA; san1Δ::kanMX4 ;ubr1Δ::hphMX* | This study |
| **DFY058** | *MATa; his3Δ1; leu2Δ0; met15Δ0; ura3Δ0:: PTEF1-stGnd1-HA; san1Δ::kanMX4 ;ubr1Δ::hphMX* | This study |
| **DFY115** | *MATa his3Δ1, leu2Δ0 met15Δ0 ura3Δ0:: PPIR3-prog-inducible synTA-Zif268 DBD; san1Δ::kanMX4* | This study |
| **DFY116** | *MATa his3Δ1, leu2Δ0 met15Δ0 ura3Δ0:: PPIR3-prog-inducible synTA-Zif268 DBD; ubr1Δ::kanMX4* | This study |
| **DFY117** | *MATa his3Δ1, leu2Δ0 met15Δ0 ura3Δ0:: PPIR3-prog-inducible synTA-Zif268 DBD; san1Δ::kanMX4; ubr1Δ::hphMX* | This study |
| **DFY118** | *MATa his3Δ1:: pZ-tGnd1-HA, leu2Δ0 met15Δ0 ura3Δ0:: PPIR3-prog-inducible synTA-Zif268 DBD; san1Δ::kanMX4* | This study |
| **DFY119** | *MATa his3Δ1:: pZ-tGnd1-3HA, leu2Δ0 met15Δ0 ura3Δ0:: PPIR3-prog-inducible synTA-Zif268 DBD; ubr1Δ::kanMX4* | This study |
| **DFY120** | *MATa his3Δ1:: pZ-tGnd1-3HA, leu2Δ0 met15Δ0 ura3Δ0:: PPIR3-prog-inducible synTA-Zif268 DBD; san1Δ::kanMX4; ubr1Δ::hphMX* | This study |
| **DFY121** | *MATa his3Δ1:: pZ-stGnd1-3HA, leu2Δ0 met15Δ0 ura3Δ0:: PPIR3-prog-inducible synTA-Zif268 DBD; san Δ::kanMX4* | This study |
| **DFY122** | *MATa his3Δ1:: pZ-stGnd1-3HA, leu2Δ0 met15Δ0 ura3Δ0:: PPIR3-prog-inducible synTA-Zif268 DBD; ubr1Δ::kanMX4* | This study |
| **DFY123** | *MATa his3Δ1:: pZ-stGnd1-3HA, leu2Δ0 met15Δ0 ura3Δ0:: PPIR3-prog-inducible synTA-Zif268 DBD; san1Δ::kanMX4; ubr1Δ::hphMX* | This study |
| **DFY192** | *MATα; his3Δ1::PPIR3-NGreen-tGnd1-HA, HIS3; leu2Δ0; met15Δ0; ura3Δ0::Nup49-mScarlet, URA3* | This study |
| **DFY193** | *MATa; his3Δ1::PPIR3-NGreen-tGnd1-HA, HIS3; leu2Δ0; met15Δ0; ura3Δ0::Nup49-mScarlet, URA3; san1Δ::kanMX4* | This study |
| **DFY194** | *MATa; his3Δ1::PPIR3-NGreen-tGnd1-HA, HIS3; leu2Δ0; met15Δ0; ura3Δ0::Nup49-mScarlet, URA3; ubr1Δ::kanMX4* | This study |
| **DFY195** | *MATa; his3Δ1::PPIR3-NGreen-tGnd1-HA, HIS3; leu2Δ0; met15Δ0; ura3Δ0::Nup49-mScarlet; san1Δ::kanMX4 ; ubr1Δ::hphMX* | This study |
| **DFY196** | *MATα; his3Δ1::PPIR3-NGreen-stGnd1-HA, HIS3; leu2Δ0; met15Δ0; ura3Δ0::Nup49-mScarlet-URA3* | This study |
| **DFY197** | *MATa; his3Δ1::PPIR3-NGreen-stGnd1-HA, HIS3; leu2Δ0; met15Δ0; ura3Δ0::Nup49-mScarlet, URA3; san1Δ::kanMX4* | This study |
| **DFY198** | *MATa; his3Δ1::PPIR3-NGreen-stGnd1-HA, HIS3; leu2Δ0; met15Δ0; ura3Δ0::Nup49-mScarlet, URA3; ubr1Δ::kanMX4* | This study |
| **DFY200** | *MATa; his3Δ1::PPIR3-NGreen-tGnd1-HA, HIS3; leu2Δ0; met15Δ0; ura3Δ0::Nup49-mScarlet; san1Δ::kanMX4 ; ubr1Δ::hphMX* | This study |
| **MBY482** | *MATα; his3Δ1; leu2Δ0; met15Δ0; ura3Δ0::PPIR3-tGnd1-HA URA3, cue5Δ::kanMX4* | This study |
| **MBY483** | *MATα; his3Δ1; leu2Δ0; met15Δ0; ura3Δ0::PPIR3-tGnd1-HA URA3, atg1Δ::kanMX4* | This study |
| **MBY484** | *MATα; his3Δ1; leu2Δ0; met15Δ0; ura3Δ0::PPIR3-tGnd1-HA URA3, atg8Δ::kanMX4* | This study |
| **MBY486** | *MATα; his3Δ1; leu2Δ0; met15Δ0; ura3Δ0::PPIR3-stGnd1-HA URA3, cue5Δ::kanMX4* | This study |
| **MBY487** | *MATα; his3Δ1; leu2Δ0; met15Δ0; ura3Δ0::PPIR3-stGnd1-HA URA3, atg1Δ::kanMX4* | This study |
| **MBY488** | *MATα; his3Δ1; leu2Δ0; met15Δ0; ura3Δ0::PPIR3-stGnd1-HA URA3, atg8Δ::kanMX4* | This study |
| **MBY501** | *his3Δ1; leu2Δ0; ura3Δ0::PPIR3-tGnd1-HA URA3; met15Δ0; GFP-ATG8* | This study |
| **MBY507** | *his3Δ1; leu2Δ0; ura3Δ0::PPIR3-tGnd1-HA URA3; met15Δ0; GFP-ATG8, atg1:: kanMX4* | This study |
| **MBY513** | *MATα; his3Δ1; leu2Δ0; met15Δ0; ura3Δ0::PPIR3-tGnd1-HA URA3* | This study |
| **MBY514** | *MATα; his3Δ1; leu2Δ0; met15Δ0; ura3Δ0::PPIR3-stGnd1-HA URA3* | This study |

| | | |
|---|---|---|
| **MPY152** | *MATα; his3Δ1; leu2Δ0; met15Δ0; ura3Δ0::PPIR3-tGnd1-HA URA3* | This study |
| **MPY153** | *MATα; his3Δ1; leu2Δ0; met15Δ0; ura3Δ0::PPIR3-stGnd1-HA URA3* | This study |
| **MPY154** | *MATα; his3Δ1; leu2Δ0; met15Δ0; ura3Δ0::PTEF1-tGnd1-HA URA3* | This study |
| **MPY155** | *MATa his3Δ1 leu2Δ0 met15Δ0 ura3Δ0::PTEF1-stGnd1-HA URA3* | This study |
| **MPY156** | *MATα; his3Δ1; leu2Δ0; met15Δ0; ura3Δ0::PPIR3-tGnd1-HA URA3; nvj1Δ::kanMX4* | This study |
| **MPY157** | *MATα; his3Δ1; leu2Δ0; met15Δ0; ura3Δ0::PPIR3-tGnd1-HA URA3; vac8Δ::kanMX4* | This study |
| **MPY158** | *MATα; his3Δ1; leu2Δ0; met15Δ0; ura3Δ0::PPIR3-stGnd1-HA URA3; nvj1Δ::kanMX4* | This study |
| **MPY159** | *MATα; his3Δ1; leu2Δ0; met15Δ0; ura3Δ0::PPIR3-stGnd1-HA URA3; vac8Δ::kanMX4* | This study |
| **MPY160** | *MATα; his3Δ1; leu2Δ0; met15Δ0; ura3Δ0::PTEF1-tGnd1-HA URA3; nvj1Δ::kanMX4* | This study |
| **MPY161** | *MATα; his3Δ1; leu2Δ0; met15Δ0; ura3Δ0::PTEF1-tGnd1-HA URA3; vac8Δ::kanMX4* | This study |
| **MPY162** | *MATα; his3Δ1; leu2Δ0; met15Δ0; ura3Δ0::PTEF1-stGnd1-HA URA3; nvj1Δ::kanMX4* | This study |
| **MPY163** | *MATα; his3Δ1; leu2Δ0; met15Δ0; ura3Δ0::PTEF1-stGnd1-HA URA3; vac8Δ::kanMX4* | This study |
| **MPY164** | *MATα; his3Δ1; leu2Δ0; met15Δ0; ura3Δ0::PPIR3-tGnd1-HA URA3; nvj1Δ::kanMX4; atg1Δ::hphMX* | This study |
| **MPY166** | *MATa/MATα; leu2-3,112; trp1-1; can1-100; ura3-1:: PPIR3-tGnd1-HA, URA3; ade2-1; his3-11,15; phi+* | This study |
| **MPY167** | *MATa/MATα; leu2-3,112; trp1-1; can1-100; ura3-1:: PPIR3-stGnd1-HA, URA3; ade2-1; his3-11,15; phi+* | This study |
| **MPY170** | *MATa/MATα; leu2-3,112; trp1-1; can1-100; ura3-1:: PPIR3-tGnd1-HA, URA3; ade2-1; his3-11,15; phi+ ; atg1:: kanMX4* | This study |

**Table 2.** Yeast strain construction

| | Construction | |
|---|---|---|
| **Yeast strain constructed** | DNA used for transformation of the starting strain/ restriction enzyme used to cut the DNA | Starting strain that was transformed with the indicated DNA |
| **DFY001** | pDF037/ AscI | BY4741 |
| **DFY002** | pDF038/ AscI | BY4741 |
| **DFY003** | pDF039/ AscI | BY4741 |
| **DFY004** | pDF040/ AscI | BY4741 |
| **DFY005** | pDF041/ AscI | BY4741 |
| **DFY006** | pDF042/ AscI | BY4741 |
| **DFY037** | pDF037/ AscI | Y04077 |
| **DFY038** | pDF038/ AscI | Y04077 |
| **DFY039** | pDF040/ AscI | Y04077 |
| **DFY040** | pDF041/ AscI | Y04077 |
| **DFY041** | pDF037/ AscI | Y04814 |
| **DFY042** | pDF038/ AscI | Y04814 |
| **DFY043** | pDF040/ AscI | Y04814 |
| **DFY044** | pDF041/ AscI | Y04814 |
| **DFY049** | pDF068/ AscI | BY4741 |
| **DFY050** | *ubr1Δ::hphMX* (PCR using pAG32) | Y04077 |
| **DFY052** | pDF069/ PmeI | DFY049 |
| **DFY053** | pDF070/ PmeI | DFY049 |
| **DFY054** | pDF071/ PmeI | DFY049 |
| **DFY055** | pDF037/ AscI | DFY050 |
| **DFY056** | pDF038/ AscI | DFY050 |
| **DFY057** | pDF040/ AscI | DFY050 |
| **DFY058** | pDF041/ AscI | DFY050 |
| **DFY115** | pDF068/ AscI | Y04077 |
| **DFY116** | pDF068/ AscI | Y04814 |
| **DFY117** | pDF068/ AscI | DFY050 |
| **DFY118** | pDF069/ PmeI | DFY115 |
| **DFY119** | pDF069/ PmeI | DFY116 |
| **DFY120** | pDF069/ PmeI | DFY117 |
| **DFY121** | pDF070/ PmeI | DFY115 |
| **DFY122** | pDF070/ PmeI | DFY116 |
| **DFY123** | pDF070/ PmeI | DFY117 |
| **DFY145** | pDF037/ AscI | MBY307 |
| **DFY146** | pDF038/ AscI | MBY307 |
| **DFY147** | pDF040/ AscI | MBY307 |
| **DFY148** | pDF041/ AscI | MBY307 |
| **DFY192** | pDF129/ PmeI, and PCR *NUP49*-mScarlet | BY4741 |
| **DFY193** | pDF129/ PmeI, and PCR *NUP49*-mScarlet | Y04077 |
| **DFY194** | pDF129/ PmeI, and PCR *NUP49*-mScarlet | Y04814 |
| **DFY195** | pDF129/ PmeI, and PCR *NUP49*-mScarlet | DFY050 |
| **DFY196** | pDF128/ PmeI, | BY4741 |

| | | |
|---|---|---|
| | and PCR *NUP49*-mScarlet | |
| **DFY197** | pDF128/ PmeI,<br>and PCR *NUP49*-mScarlet | Y04077 |
| **DFY198** | pDF128/ PmeI,<br>and PCR *NUP49*-mScarlet | Y04814 |
| **DFY200** | pDF128/ PmeI,<br>and PCR *NUP49*-mScarlet | DFY050 |
| **MBY482** | pDF037/ AscI | Y01818 |
| **MBY483** | pDF037/ AscI | Y04547 |
| **MBY484** | pDF037/ AscI | Y03104 |
| **MBY486** | pDF038/ AscI | Y01818 |
| **MBY487** | pDF038/ AscI | Y04547 |
| **MBY488** | pDF038/ AscI | Y03104 |
| **MBY513** | pDF037/ AscI | BY4741 |
| **MBY514** | pDF038/ AscI | BY4741 |
| **MPY152** | pDF037/ AscI | BY4741 |
| **MPY153** | pDF038/ AscI | BY4741 |
| **MPY154** | pDF040/ AscI | BY4741 |
| **MPY155** | pDF041/ AscI | BY4741 |
| **MPY156** | pDF037/ AscI | Y02889 |
| **MPY157** | pDF037/ AscI | Y00253 |
| **MPY158** | pDF038/ AscI | Y02889 |
| **MPY159** | pDF038/ AscI | Y00253 |
| **MPY160** | pDF040/ AscI | Y02889 |
| **MPY161** | pDF040/ AscI | Y00253 |
| **MPY162** | pDF041/ AscI | Y02889 |
| **MPY163** | pDF041/ AscI | Y00253 |
| **MPY164** | *atg1Δ::hphMX*<br>(PCR using pAG32) | MPY156 |
| **MPY166** | pDF037 / AscI | W303 |
| **MPY167** | pDF038/ AscI | W303 |
| **MPY170** | *atg1Δ::hphMX*<br>(PCR using pAG32) | MPY166 |

**Table 3.** Primers used in this study

| Primer | Sequence 5' → 3' | Purpose |
|---|---|---|
| **prDF003** | GTTGTTGGTACCGGTTTCTAAAATGTGCAACC | Cloning *PIR3* promoter |
| **prDF004** | TGTTGTTCTCGAGACTTATAAACAGTACTTGTTTTATGAG | Cloning *PIR3* promoter |
| **prDF050** | GTGTTCTCGAGGTTGCCTAGGGACTTATAAACAGTACTTGttttatg | Cloning *PPIR3* (AvrII)-s/tGnd1 constructs |
| **prDF051** | AGGAACCTAGGaatggtctctaagggtgaag | Cloning mNeonGreen-tagged s/tGnd1 |
| **prDF052** | AGGAACTCGAGACCAGAAGAACCACCACCACCAGAACCcttgtacaattcgtccatacc | Cloning mNeonGreen-tagged s/tGnd1 |
| **prKZ129** | ACTGTTTATGGATATCGCTGAGAGAATCGCCGTGTTACATCAAAAAACGAAAACACTGGCATCATTGAGCATAATCGGTGACGGTGCTGGT | C-terminal tagging of Nup49 with mScarlet (Nup49-mScarlet) |
| **prKZ130** | TATCACACTATTAGCAATGACTTTCACATTTTAAAGGAAAATAATAGTATTAAAAATTAAACAGTGACGAAGGAAGGCAGCAGTATAGCGACCAGCAT | C-terminal tagging of Nup49 with mScarlet (Nup49-mScarlet) |
| **prMB601** | GTTGTTGGTACCACACACCATAGCTTCA | Construction of pMB151 (amplification of *PTEF1*) |
| **prMB602** | GTTGTTCTCGAGTTGTAATTAAAACTTAGATTAG | Construction of pMB151 (amplification of *PTEF1*) |
| **prMB603** | GTTGTTTCTAGATCATGTAATTAGTTATGTCAC | Construction of pMB152 (amplification of *TCYC1*) |
| **prMB604** | GTTGTTGAGCTCGCAAATTAAAGCCTTCGAGCG | Construction of pMB152 (amplification of *TCYC1*) |
| **prMB683** | ATTTACATCAATAAGAAATCTCATAAAACAAGTACTGTTTATAAGTCTCGAGATGTCTGCTGATTTCGGTTTG | Cloning *GND1*, *tGND1* and *stGND1* |
| **prMB684** | ACGTCATAGGGATAGCCCGCATAGTCAGGAACATCGTATGGGTATCTAGaACCAGCTTGGTATGTAGAGGAAG | Cloning *GND1* |
| **prMB685** | TTCTAGATACCCATACGATGTTCCTGACTATGCGG | Cloning HA-epitope tag |
| **prMB686** | AACGTCATATGGATAGGATCCTGCATAGTCCGGGACGTCATAGGGATAGCCCGCATAGTCAGGAACATCGTATGG | Cloning HA-epitope tag |
| **prMB687** | TGCAGGATCCTATCCATATGACGTTCCAGATTACGCTTCTAGATGATGAACTAGTTCTAGATCATGTAATTAGTT | Cloning HA-epitope tag |
| **prMB688** | GAGGGCGTGAATGTAAGCGTGACATAACTAATTACATGATCTAGAACTAG | Cloning HA-epitope tag |
| **prMB689** | ACGTCATAGGGATAGCCCGCATAGTCAGGAACATCGTATGGGTATCTAGaACCTGGCCAAGCTTCTTCAGAAC | Cloning stGND1 |
| **prMB690** | ACGTCATAGGGATAGCCCGCATAGTCAGGAACATCGTATGGGTATCTAGaACCTCTAATGATACAACCACCTC | Cloning tGND1 |
| **prMB718f** | GTTGTTctcgagaataaaGCACGAactagtGGTtCTAGATACCCATACGATGTTCCTGACTATGCGTGATAAccgcggAACAAC | Construction of plasmid with 3HA for protein tagging |

| | | |
|---|---|---|
| **prMB718r** | GTTGTTccgcggTTATCACGCATAGTCAGGAACATCGTATGG GTATCTAGaACCactagtTCGTGCtttattctcgagAACAAC | Construction of plasmid with 3HA for protein tagging |
| **prMB753** | GTTGTTGAATTCGCAGGCTAGCAATAAAATG | ZF/GAL4-DBD-progest.-TA |
| **prMB755** | GTTGTTACTAGTCGTATATAATTTAGCTATTTGCTTA | ZF/GAL4-DBD-progest.-TA |

**Table 4.** Plasmids used in this study

| Plasmid | Description | Reference |
|---|---|---|
| **pAG32** | hphMX4, pFA6 | (69) |
| **pDF037** | *PPIR3*-tGnd1-HA, *URA3*, integrative (based on pRG206MX) | This study |
| **pDF038** | *PPIR3*-stGnd1-HA, *URA3*, integrative (based on pRG206MX) | This study |
| **pDF039** | *PPIR3*-Gnd1-HA, *URA3*, integrative (based on pRG206MX) | This study |
| **pDF040** | *PTEF1*-tGnd1-HA, *URA3*, integrative (based on pRG206MX) | This study |
| **pDF041** | *PTEF1*-stGnd1-HA, *URA3*, integrative (based on pRG206MX) | This study |
| **pDF042** | *PTEF1*-Gnd1-HA, *URA3*, integrative (based on pRG206MX) | This study |
| **pDF068** | *PPIR3*- prog-inducible synTA-Zif268 DBD, *URA3*, integrative (based on pRG206MX) | This study |
| **pDF069** | pZ-tGnd1-HA, *HIS3*, integrative (based on pHES836) | This study |
| **pDF070** | pZ-stGnd1-HA, *HIS3*, integrative (based on pHES836) | This study |
| **pDF071** | pZ-Gnd1-HA, *HIS3*, integrative (based on pHES836) | This study |
| **pDF072** | *PPIR3* (AvrII) - tGND1-3HA, URA3, integrative (based on pRG206MX) | This study |
| **pDF073** | *PPIR3* (AvrII) - stGND1-3HA, URA3, integrative (based on pRG206MX) | This study |
| **pDF126** | *PPIR3*-stGnd1-HA, *HIS3*, integrative (based on pMB278) | This study |
| **pDF127** | *PPIR3*-tGnd1-HA, *HIS3*, integrative (based on pMB278) | This study |
| **pDF128** | *PPIR3*-NeonGreen-stGnd1-HA *HIS3*, integrative (based on pMB278) | This study |
| **pDF129** | *PPIR3*-NeonGreen-tGnd1-HA *HIS3*, integrative (based on pMB278) | This study |
| **pFA6a-link-ymNeongreen-SpHis5** | linker-ymNeonGreen (pFA6) | (43) gift from Bas Teusink (Addgene #125704) |
| **pFA6a-link-ymScarletI-URA3** | linker-ymScarlet (pFA6) | (43) gift from Bas Teusink (Addgene #168055) |
| **pMB151** | *PTEF1*, *URA3,* CEN (based on pRG216) | This study |
| **pMB152** | *PTEF1*- multicloning site -TCYC1, *URA3,* CEN (based on pRG216) | This study |
| **pMB211** | *PPIR3*-tGnd1-HA, *URA3*, CEN (based on pRG216) | This study |
| **pMB212** | *PPIR3*-stGnd1-HA, *URA3*, CEN (based on pRG216) | This study |
| **pMB213** | *PPIR3*-Gnd1-HA, *URA3*, CEN (based on pRG216) | This study |
| **pMB214** | *PTEF1*-tGnd1-HA, *URA3*, CEN (based on pRG216) | This study |
| **pMB215** | *PTEF1*-stGnd1-HA, *URA3*, CEN (based on pRG216) | This study |
| **pMB216** | *PTEF1*-Gnd1-HA, *URA3*, CEN (based on pRG216) | This study |
| **pMB272** | *PTEF1*-multicloning site -TCYC1, *URA3,* integrative (based on pRG206MX) | This study |
| **pMB274** | *PTEF1*-progesterone-induced ZifDBD-TA-TCYC1, *URA3,* integrative (based on pRG206MX) | This study |
| **pMB278** | pZ-multicloning site-HA tag, *HIS3*, integrative (based on pHES836) | This study |
| **pHES830** | *Zif268 DBD - hPR LBD - MSN2 AD, URA3,* integrative | (70) gift from Hana El-Samad (Addgene #87944) |
| **pHES836** | pZ-mKate2, *HIS3* | (70) gift from Hana El-Samad (Addgene #89195) |
| **pRG206MX** | *URA3,* integrative | (71) gift from Joerg Stelling (Addgene #64536) |
| **pRG216** | *URA3,* CEN | (71) gift from Joerg Stelling (Addgene #64536) |
| **pXP732** | *CYC1* terminator insert, *URA3, CEN* | (72) gift from Nancy DaSilva (Addgene #46058) |

**Table 5.** Plasmid construction

| Plasmid | Description of the construction |
|---|---|
| **pDF037**<br>**pDF038**<br>**pDF039** | To obtain integrative plasmids, *PPIR3*-tGND1-HA, *PPIR3*-stGND1-HA and *PPIR3*-*GND1*-HA constructs from pMB211, pMB212 and pMB213, respectively, were cut with KpnI and SpeI and ligated into pMB272 vector. |
| **pDF040**<br>**pDF041**<br>**pDF042** | To obtain integrative plasmids, tGND1-HA, stGND1-HA and GND1-HA constructs from CEN-plasmids pMB214, pMB215 and pMB216, respectively, were cut with XhoI and SpeI and the inserts were ligated into pMB272 vector, yielding pDF040, pDF041 and pDF042. |
| **pDF067** | *TEF1*-promoter in pMB272 was exchanged to *PIR3*-promoter by ligating *PIR3*-promoter sequence into KpnI and XhoI-cut pMB272, yielding pDF067. *PIR3*-gene promoter had been PCR amplified from genomic DNA of wild-type BY4741 with primers prDF003 and prDF004. |
| **pDF068** | To place synthetic transcription factor (prog-inducible synTA-Zif268 DBD) under the control of *PIR3*-promoter, the insert was cut from pMB274 with EcoRI and SpeI, and ligated into similarly cut pDF067 vector containing *PIR3*-promoter. |
| **pDF069**<br>**pDF070**<br>**pDF071** | To place tGND1-HA, stGND1-HA and GND1-HA constructs under the control of Z-promoter, pDF040, pDF041 or pDF042 plasmids were cut with XhoI/ SacI and the inserts were ligated into pMB278, yielding pDF069, pDF070 and pDF071, respectively. |
| **pDF072** | *PPIR3* containing an AvrII restriction site was amplified with primers pDF003 and 050 from pDF037, cut with XhoI and KpnI and ligated with similarly cut pDF037. |
| **pDF073** | *PPIR3* containing an AvrII restriction site was amplified from pDF037 using primers pDF003 and 050, cut with XhoI / KpnI and ligated with similarly cut pDF038. |
| **pDF126**<br>**pDF127** | *PPIR3*-stGnd1-HA and *PPIR3*-tGnd1-HA inserts were cut from plasmids pDF073 (stGnd1) and pDF072 (tGnd1), respectively, with KpnI/ SacI, and ligated with similarly cut pMB278. |
| **pDF128**<br>**pDF129** | To insert mNeonGreen at the N-terminus of stGnd1-HA and tGnd1-HA, mNeonGreen was amplified from pFA6a-link-ymNeongreen-SpHis5 using primers prDF051 and prDF052, cut with AvrII / XhoI and ligated with similarly cut pDF126 and pDF127, yielding pDF128 or pDF129, respectively. |
| **pMB151** | PCR product encompassing *TEF1*-promoter (obtained by primers prMB601, and prMB602 and genomic DNA of wild-type strain BY4742 as a template) was cut with KpnI and XhoI, and ligated with similarly cut pRG216 vector. |
| **pMB152** | PCR product encompassing *CYC1*- terminator (obtained by primers prMB603 and prMB604, and pXP732 as a template) was cut with XbaI and SacI and ligated with similarly cut pMB151 vector. |
| **pMB211**<br>**pMB212**<br>**pMB213** | Cloning of plasmids pMB211-213 was done in several steps. To clone pMB211, first, tGND1 missing codons for the last 121 amino acids was amplified by PCR from genomic DNA of BY4741 using primers prMB683 and prMB690, yielding DNA fragment "A-211". To clone pMB212, first, stGND1 (missing codons for the last 339 amino acids) was cloned in several steps: was amplified by PCR from genomic DNA of BY4741 using primers prMB683 and prMB689, yielding DNA fragment "A-212". To clone pMB213, first, *GND1* was amplified by PCR from genomic DNA of BY4741 using primers prMB683 and prMB684, yielding DNA fragment "A-213". |

| | |
|---|---|
| | Second, 3HA cassette was assembled by annealing of oligonucleotides prMB685, 686, 687, 688 and amplified by PCR, yielding DNA fragment "B".<br>Third, DNA fragment "C" was XhoI/ SpeI-cut vector containing *PPIR3* and *TCYC1,* and a *URA3* selection marker.<br>Finally, plasmids pMB211, pMB212 and pMB213 were created using homologous recombination, by co-transforming a Ura- yeast strain with the DNA fragments "A", "B" and "C", followed by selection for Ura+ colonies. |
| **pMB214**<br>**pMB215**<br>**pMB216** | To place tGND1-HA, stGND1-HA or GND1-HA construct under the control of *TEF1*-promoter, the constructs were cut from pMB211, pMB212 or pMB213, respectively, by XhoI/ SpeI, and ligated with similarly cut pMB152 vector. |
| **pMB272** | To obtain integrative *URA3* plasmid carrying PTEF1-linker-TCYC1 construct, pMB152 was cut with EagI and this insert was ligated into EagI-cut pRG206MX vector. |
| **pMB274** | PCR product amplified from pHES830 with prMB753 and prMB755 was cut with EcoRI and SpeI and ligated into pMB272 vector. |
| **pMB278** | 3HA was amplified from pMB211 with primers prMB718-f and prMB718-r, cut with XhoI and SacII, and ligated into pHES836 vector. |
